# Modeling movie-evoked human brain activity using motion-energy and space-time vision transformer features

**DOI:** 10.1101/2021.08.22.457251

**Authors:** Shinji Nishimoto

## Abstract

In this paper, the process of building a model for predicting human brain activity under video viewing conditions was described as a part of an entry into the Algonauts Project 2021 Challenge. The model was designed to predict brain activity measured using functional MRI (fMRI) by weighted linear summations of the spatiotemporal visual features that appear in the video stimuli (video features). Two types of video features were used: (1) motion-energy features designed based on neurophysiological findings, and (2) features derived from a space-time vision transformer (TimeSformer). To utilize the features of various video domains, the features of the TimeSformer models pre-trained using several different movie sets were combined. Through these model building and validation processes, results showed that there is a certain correspondence between the hierarchical representation of the TimeSformer model and the hierarchical representation of the visual system in the brain. The motion-energy features are effective in predicting brain activity in the early visual areas, while TimeSformer-derived features are effective in higher-order visual areas, and a hybrid model that uses motion energy and TimeSformer features is effective for predicting whole brain activity.

## Background and Aims

Building predictive models of the brain is a fundamental goal in systems neuroscience. Previous studies have modeled brain activity evoked by naturalistic movie stimuli using various visual features, including motion-energy features derived from neurophysiological findings (Watson and Ahumada 1985; Nishimoto et al., 2011) and convolutional neural network (CNN)-based features derived from the context of computational image classification tasks (Tran et al., 2015; Guclu and van Gerven 2017). In recent years, vision transformers (ViT; Dosovitskiy et al., 2020) have become a popular algorithm for image classification, but relatively little is known about how the visual features learned by these models correspond to brain representations, especially brain activity evoked by movie stimuli. This study used the TimeSformer (Bertasius et al., 2021), a space-time vision transformer that handles movie inputs, to examine the possibility of using its internal representations for predicting movie-evoked brain activity.

## Methods

Voxel-wise encoding models (Naselaris et al., 2011) were built to predict brain activity evoked by movie stimuli. The models consist of a visual feature extraction stage followed by a weighted linear summation. Different classes of features derived from either TimeSformer (Bertasius et al., 2021) or motion energy (Watson and Ahumada 1985) models were examined.

### TimeSformer features

To extract video features derived from the vision transformer, TimeSformer (Bertasius et al., 2021) models that were pre-trained using various movie datasets were used. The model contained 12 layers of video vision transformer blocks. The output of each block was used as the feature representation for each layer. To reduce the total number of features, the features over the spatial encoding dimension were integrated, thereby obtaining a total of 768 channel features for each of the 12 layers.

To obtain features for a variety of video contents, the models that were pre-trained using the following video sets were used: Kinetics-400 (K400; Carreira and Zisserman 2017), Kinetics-600 (K600; Carreira et al., 2018), Something-Something-V2 (SSv2; Goyal et al., 2017), and HowTo100M (Miech et al., 2019). The TimeSformer code and pre-trained models were obtained from the following publicly available sources: https://github.com/facebookresearch/TimeSformer.

### Motion-energy features

To use visual features derived from neurophysiological findings, motion-energy filters that extract visually localized motion components in the video were used (Watson and Ahumada 1985; Nishimoto et al, 2011). The motion-energy filters mimic the response of neurons in the early visual cortex, and each filter detects motion components of a specific direction and speed that appear at a specific location in the visual field. More specifically, the video was first transformed into Commission Internationale de l’Eclairage (CIE) LCh color space, and only the luminance component (L) was used for the following analysis, unless otherwise noted. A space-time 3-dimensional Gabor filter was used to detect each motion component, and the phase-independent motion component was quantified by the squared sum of the output of Gabor filters with a quadrature pair. A total of 2,139 channels of motion-energy filters were used to detect motions at each position in the visual field covering various motion directions (eight directions), spatial frequencies (0, 1.5, 3, 6, 12, and 24 cycles/visual field), and temporal frequencies (0, 4, and 8 Hz). In some analyses, a high-resolution model (6,555 channels) with a maximum spatial frequency component of 32 cycles/visual field was used. To calculate the motion-energy features, we used the following publicly available code: https://github.com/gallantlab/motion_energy_matlab

### Data

Data published by the Algonauts Project 2021 (Cichy et al., 2021) was used. Specifically, the data used for training encoding models consisted of 1000 3-second movie clips and the movie-evoked brain activity measured using functional magnetic resonance imaging (fMRI) for 10 participants. Using the trained models, brain activity prediction for separate test movie sets were performed (102 of 3-second movie clips), and the prediction results were submitted to the Algonauts Project 2021 Challenge.

### Model regression

We built voxel-wise encoding models to predict movie-evoked brain activity (Naselaris et al., 2011; Nishimoto et al., 2011). Given the visual features S, feature-wise weight matrix W, and noise e, the model for predicting brain activity R was

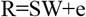

To obtain the optimal W, an L2-regularized linear regression (ridge regression) was applied. To optimize the regularization parameters, the hold-out method was used, in which 90% of the training data were used for regression and the remaining 10% were used for validation. For multivariate ridge regression, the code ridgemulti published at https://osf.io/ea2jc/ (Nakai and Nishimoto 2020) was used.

## Results

### TimeSformer: multilayer representation

To examine the effectiveness of video features obtained from transformer hierarchical representation on brain activity predictions, voxel-wise encoding models were constructed using the following features derived from a TimeSformer model: (1) outputs of each layer (i.e., layers 1-12) and (2) the sum of the outputs of all layers. The TimeSformer model that was pretrained using the K400 movie set was used. The prediction accuracy (correlation coefficient between the predicted and measured brain activity) was then quantified on the validation data and summarized by averaging the accuracy for each region of interest (ROI) (Figure 1).

**Fig. 1.**
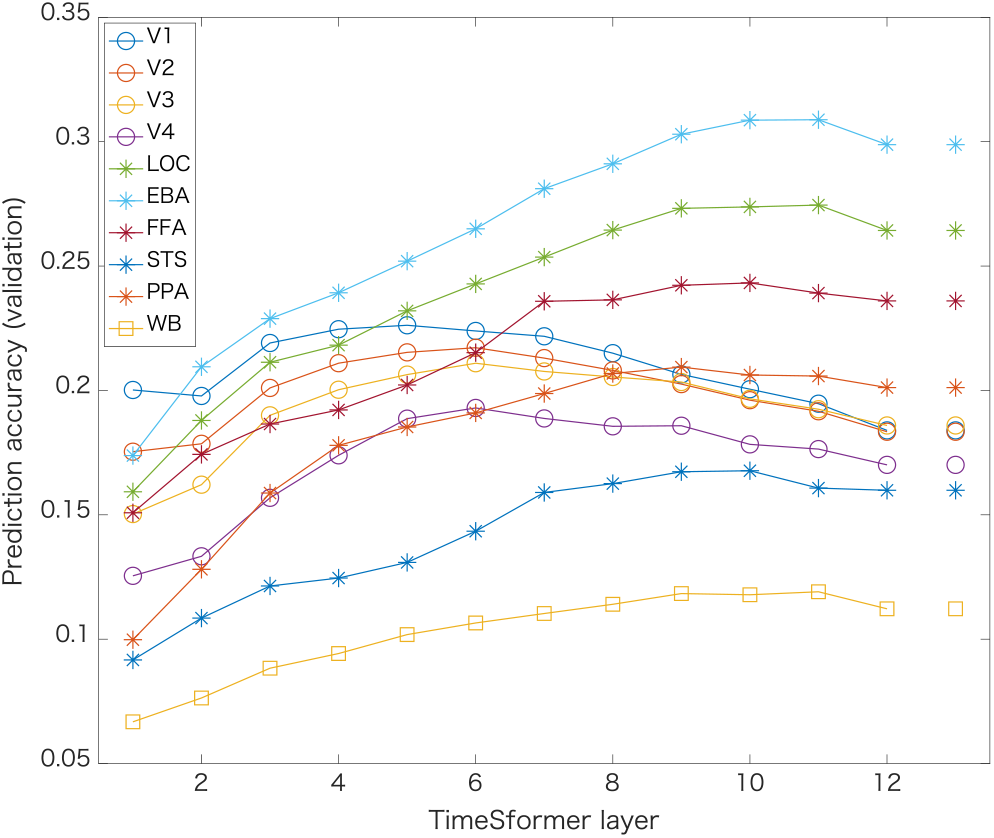
Prediction accuracy of encoding models using the output of each layer (block) of the TimeSformer model. Plots show the model prediction accuracy (correlation coefficient) of the validation data for the model using the features of each TimeSformer layer (1-12) and the sum of them (rightmost). The plots with colors and icons indicate the prediction accuracy for each ROI (see legend; LOC: lateral occipital complex, EBA: extrastriate body area, FFA: fusiform face area, STS: superior-temporal sulcus, PPA: parahippocampal place area, WB: whole brain.)

The model using the features of the lower layers (layers 4-6) was found to be effective in the early visual cortex (V1-V4), while the model using the features of the higher layers (layers 9-11) was effective in the higher visual cortex such as the FFA. For the higher visual areas and the whole brain, the model using the sum of the outputs of all layers was found to be roughly representative of the accuracy for each ROI.

### TimeSformer: movie set dependency

Each TimeSformer model was pre-trained on a specific set of videos (e.g., human actions), and these cover a portion of the total video features that humans recognize. By combining video features derived from multiple video sets, the total feature set cover more comprehensive video features, and thus might explain a larger variance of movie-evoked brain activity. To test this hypothesis, the prediction accuracy of encoding models using features (the sum of all layers) of the TimeSformer models that were pre-trained on four video sets (K400, K600, SSv2, and HowTo100M) were quantified, as well as an encoding model concatenating the features of these four TimeSformer models (Table 1).

**Table 1.**
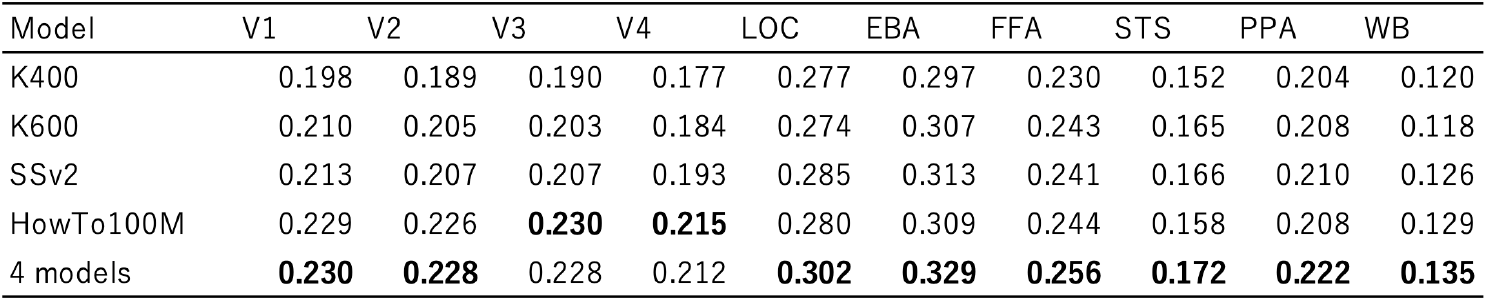
Comparison of the prediction performance of encoding models using TimeSformer feature pre-trained on four different movie sets. The prediction accuracy of models that combined features from the four video sets was also examined (bottommost).

| Model | V1 | V2 | V3 | V4 | LOC | EBA | FFA | STS | PPA | WB |
| --- | --- | --- | --- | --- | --- | --- | --- | --- | --- | --- |
| K400 | 0.198 | 0.189 | 0.190 | 0.177 | 0.277 | 0.297 | 0.230 | 0.152 | 0.204 | 0.120 |
| K600 | 0.210 | 0.205 | 0.203 | 0.184 | 0.274 | 0.307 | 0.243 | 0.165 | 0.208 | 0.118 |
| SSv2 | 0.213 | 0.207 | 0.207 | 0.193 | 0.285 | 0.313 | 0.241 | 0.166 | 0.210 | 0.126 |
| HowTo100M | 0.229 | 0.226 | **0.230** | **0.215** | 0.280 | 0.309 | 0.244 | 0.158 | 0.208 | 0.129 |
| 4 models | **0.230** | **0.228** | 0.228 | 0.212 | **0.302** | **0.329** | **0.256** | **0.172** | **0.222** | **0.135** |

For most ROIs, the best prediction performance was obtained when the features from the four models were concatenated. For V3 and V4, the model using HowTo100M-derived features outperformed the concatenated model.

### Comparing motion-energy and TimeSformer representations

To compare the effectiveness of traditional motion-energy features and transformer-derived features, we compared brain activity prediction models using feature representations in the following three conditions: (1) motion-energy features, (2) TimeSformer features, and (3) concatenation of (1) and (2). The modeling accuracy was examined using validation data, as described above, and summarized in Table 2.

**Table 2.**
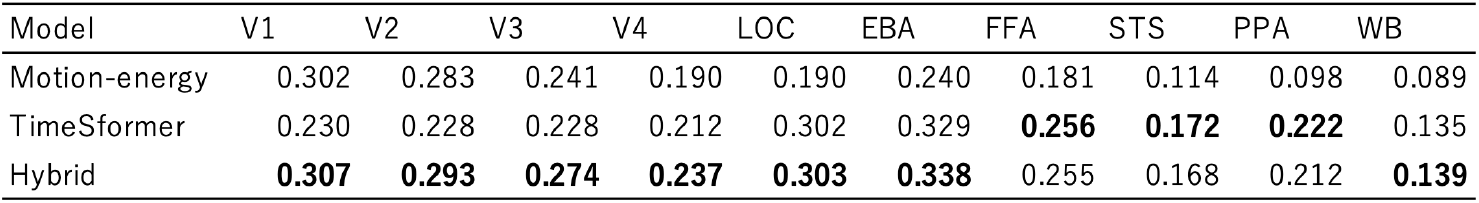
Comparison of prediction performance of encoding models using motion-energy and TimeSformer features. The values represent the average prediction accuracy (correlation coefficient) of validation data in each ROI.

The results showed that, in the early visual cortex, the model using motion-energy features showed high prediction accuracy, while in the higher visual cortex, the TimeSformer feature-based model outperformed the motion-energy model. The hybrid model that concatenated the two feature sets showed the best performance in many ROIs.

### Final Model

In the challenge phase of Algonauts Challenge 2021, the optimal feature sets (see below) that gave the best prediction accuracy (validation data) for each ROI were selected, and the final encoding models were built using all training data. The predicted activity for the test movie data was calculated and submitted to the challenge website. The following feature sets for the finalizing process were used: Motion-energy model (low resolution), Motion-energy model (high resolution), Motion-energy model (low resolution using the chromaticity channel), TimeSformer feature model (multiple video-set models concatenated)

## Discussion

In this study, the efficacy of vision transformer (ViT)-derived features in predicting movie-evoked brain activity was examined. The hierarchical internal representation of the TimeSformer model was found to correspond to the hierarchical representation in the visual system. In addition, the model using the TimeSformer features outperformed the conventional motion-energy features, except for the early visual cortex. These results suggest that ViT-based models are effective in predicting and interpreting brain activity in video viewing.

Owing to time constraints, many of the model examination processes in this study was simplified. The following possibilities can be considered for future improvements: (1) use the feature representation of the best layer for each ROI instead of the sum of all TimeSformer layers, (2) use information on position encoding, and (3) combine more diverse features, including CNN and other vision transformer models.

Although the main purpose of this study was to evaluate the accuracy of brain activity prediction for participation in the prediction challenge, it is important from a neuroscience perspective to further analyze the obtained encoding models to reveal what kind of representation the models have acquired. ViT-based models have properties that are relevant to perceptual and cognitive brain functions, such as the representation of selective attention and long-term temporal dependencies, and methods for interpreting internal transformer representations have been developed (e.g., Abnar and Zuidema 2020). Future developments are expected in terms of contrastive representational interpretation between the brain and machine learning.

## Acknowledgements

We thank the organizers of the Algonauts Project 2021 Challenge for providing this interesting opportunity. We also thank Dr. Yu Takagi for introducing this challenge. The work presented here was supported by KAKENHI JP18H05522, JST CREST JPMJCR18A5, and ERATO JPMJER1801.

## Notes

### Competing Interest Statement

The authors have declared no competing interest.

